# The repetitive genome of the *Ixodes ricinus* tick reveals transposable elements have driven genome evolution in ticks

**DOI:** 10.1101/2024.03.13.584159

**Authors:** Isobel Ronai, Rodrigo de Paula Baptista, Nicole S. Paulat, Julia C. Frederick, Tal Azagi, Julian W. Bakker, Katie C. Dillon, Hein Sprong, David A. Ray, Travis C. Glenn

## Abstract

Ticks are obligate blood-feeding parasites associated with a huge diversity of diseases globally. The hard tick *Ixodes ricinus* is the key vector of Lyme borreliosis and tick-borne encephalitis in Western Eurasia. *Ixodes* ticks have large and repetitive genomes that are not yet well characterized. Here we generate two high-quality *I*. *ricinus* genome assemblies, with haploid genome sizes of approximately 2.15 Gbp. We find transposable elements comprise at least 69% of the two *I. ricinus* genomes, amongst the highest proportions found in animals. The transposable elements in ticks are highly diverse and novel, so we constructed a repeat library for ticks using our *I*. *ricinus* genomes and the genome of *I*. *scapularis*, another major tick vector of Lyme borreliosis. To understand the impact of transposable elements on tick genomes we compared their accumulation in the two *Ixodes* sister species. We find transposable elements in these two species to be drivers of genome evolution in ticks. The *I*. *ricinus* genome assemblies and our tick repeat library will be valuable resources for biological insights into this important ectoparasite. Our findings highlight that further research into the impact of transposable elements on the genomes of blood-feeding parasites is required.

## INTRODUCTION

Ticks in the genus *Ixodes* are important global parasites. In Western Eurasia, the hard tick *Ixodes ricinus* is associated with the major human diseases Lyme borreliosis and tick-borne encephalitis^1,2^. This tick species is also associated with a diversity of emerging diseases, including those caused by the pathogens *Anaplasma phagocytophilum*, *Borrelia miyamotoi*, *Neoehrlichia mikurensis*, *Spiroplasma ixodetis*, *Rickettsia helvetica, Rickettsia monacencis* and *Babesia* species^3^. These diseases are often also of veterinary relevance, for both livestock and pets^4–6^. Furthermore, the bite of *I*. *ricinus* can cause other types of diseases, such as alpha-gal syndrome (red meat allergy)^7,8^.

The threat of tick-borne diseases associated with *I*. *ricinus* is escalating^9,10^. For example, the incidence of Lyme borreliosis and tick-borne encephalitis has increased in several European countries and is appearing in new countries^11–15^. This increase in tick-borne diseases in Europe is likely caused by higher numbers of *I. ricinus* and the spread of *I. ricinus* populations^10^. The habitat expansion of *I. ricinus* is due to reforestation and increased host availability^16^. In addition, climate change is expanding the range of *I. ricinus* at its extremes of altitude in central Europe, and extremes of latitude in Scandinavia^15,17^. New tick control strategies and vaccines are therefore urgently needed, but have been impeded by our lack of knowledge of the fundamental biology of *I. ricinus*^18^.

While *I. ricinus* is of medical and veterinary importance, we have only a preliminary understanding of its genome. The current *I. ricinus* genome assembly^19^ is partial (Table 1), covering only a quarter of the expected size relative to the genome assemblies for the other major Lyme borreliosis tick vector and sister species, *I. scapularis*^20,21^. This lack of tick genome assemblies impedes comparative analyses of major tick vectors.

**Table 1.**
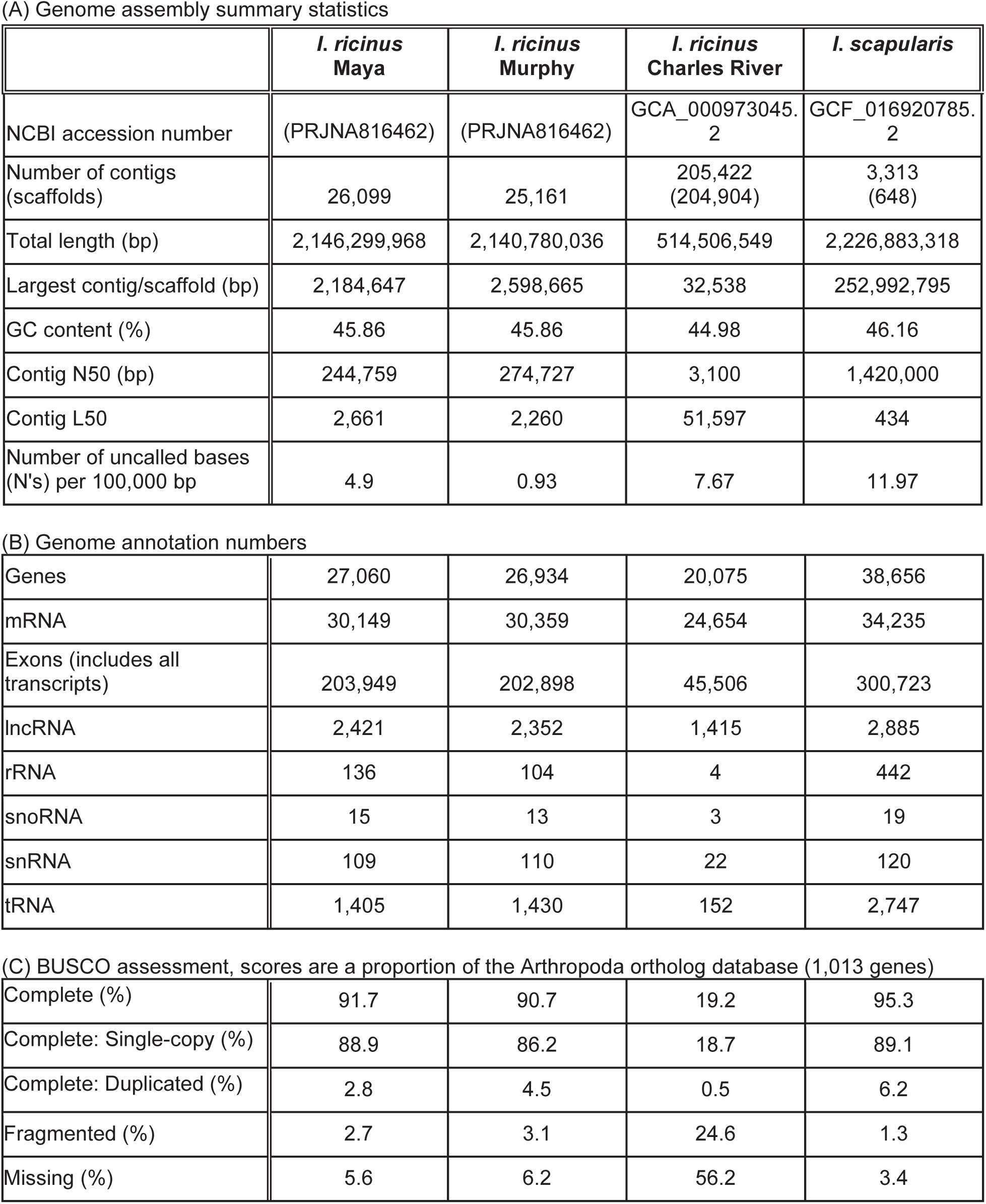
Genomes of *Ixodes ricinus* Maya and Murphy. Comparison with the *I. ricinus* partial genome assembly^19^ and *I. scapularis* reference genome assembly^20^.

(A) Genome assembly summary statistics
|  | *I. ricinus*<br>Maya | *I. ricinus*<br>Murphy | *I. ricinus*<br>Charles River | *I. scapularis* |
| --- | --- | --- | --- | --- |
| NCBI accession number | (PRJNA816462) | (PRJNA816462) | GCA_000973045.<br>2 | GCF_016920785.<br>2 |
| Number of contigs<br>(scaffolds) | 26,099 | 25,161 | 205,422<br>(204,904) | 3,313<br>(648) |
| Total length (bp) | 2,146,299,968 | 2,140,780,036 | 514,506,549 | 2,226,883,318 |
| Largest contig/scaffold (bp) | 2,184,647 | 2,598,665 | 32,538 | 252,992,795 |
| GC content (%) | 45.86 | 45.86 | 44.98 | 46.16 |
| Contig N50 (bp) | 244,759 | 274,727 | 3,100 | 1,420,000 |
| Contig L50 | 2,661 | 2,260 | 51,597 | 434 |
| Number of uncalled bases<br>(N's) per 100,000 bp | 4.9 | 0.93 | 7.67 | 11.97 |
| (B) Genome annotation numbers |  |  |  |  |
| Genes | 27,060 | 26,934 | 20,075 | 38,656 |
| mRNA | 30,149 | 30,359 | 24,654 | 34,235 |
| Exons (includes all<br>transcripts) | 203,949 | 202,898 | 45,506 | 300,723 |
| lncRNA | 2,421 | 2,352 | 1,415 | 2,885 |
| rRNA | 136 | 104 | 4 | 442 |
| snoRNA | 15 | 13 | 3 | 19 |
| snRNA | 109 | 110 | 22 | 120 |
| tRNA | 1,405 | 1,430 | 152 | 2,747 |
| (C) BUSCO assessment, scores are a proportion of the Arthropoda ortholog database (1,013 genes) |  |  |  |  |
| Complete (%) | 91.7 | 90.7 | 19.2 | 95.3 |
| Complete: Single-copy (%) | 88.9 | 86.2 | 18.7 | 89.1 |
| Complete: Duplicated (%) | 2.8 | 4.5 | 0.5 | 6.2 |
| Fragmented (%) | 2.7 | 3.1 | 24.6 | 1.3 |
| Missing (%) | 5.6 | 6.2 | 56.2 | 3.4 |

The genomes of ticks have been challenging to assemble due in part to their large genome sizes, abundant biological genome contamination and genome complexity^20–26^. In particular, a large proportion of tick genomes are repetitive, with repeats comprising more than two-thirds of the genome in *Ixodes* species^20,24,27^. However, the major component of these repetitive regions, the transposable elements (TEs), remain largely undescribed in ticks^22^. For example, we previously identified 46% of the TEs in the *I. scapularis* genome as unknown^20^. In addition, the current genome assemblies for ticks have used pools of ticks^23^ or focused on producing a consensus sequence from a single individual^20,24^, independent genome assemblies from individual ticks have not yet been generated. The generation of genome assemblies from multiple individual ticks would enable intraspecies comparisons and improve our understanding of tick biology and genomics. Here we assemble genomes for two individual field-caught *I. ricinus* females (Maya and Murphy) using a combination of long-read Oxford Nanopore sequencing and Illumina short-reads. We annotate these assemblies and characterize the novel TEs present in ticks. These two new *I. ricinus* genome assemblies enable comparisons at both the intraspecies and interspecies levels.

## RESULTS

### Two high-quality *I*. *ricinus* genome assemblies

To obtain *I*. *ricinus* genome assemblies with minimal biological genome contamination we conducted stringent filtering of the sequencing reads from *I*. *ricinus* Maya and *I*. *ricinus* Murphy. From the total number of reads, 78.6% of reads for *I*. *ricinus* Maya and 80.2% for Murphy mapped to *Ixodes* genomes (Figure S1). In addition, 10.4% of the total reads for Maya and 17.6% for Murphy were taxonomically unclassified (Figure S1) and were therefore retained as they could potentially be from *I*. *ricinus*. Both *I*. *ricinus* Maya and Murphy included biological contamination from the host blood they had been fed (0.8% and 1.3%, respectively; Figure S1) and microbiota (10.2% and 0.8%, respectively; Figure S1). The host contamination was filtered out from the reads and the microbiota contamination filtered out from the genome assemblies.

We obtained high-quality genome assemblies for *I. ricinus* Maya and Murphy. The starting genome sizes were 3.3 Gb and after purging the haplotigs, observed a haploid genome size of 2.146 Gbp for Maya and 2.140 Gbp for Murphy (Table 1A). Both Maya and Murphy have similar GC content distribution and no sequence gaps. These genome assemblies are the first released tick assemblies that use Oxford Nanopore sequencing. While our Oxford Nanopore based genome assemblies generated more contigs than the PacBio based *I. scapularis* reference genome assembly (Table 1A), these genome assemblies have comparable BUSCO completeness (Table 1C). Overall, the *I. ricinus* Maya genome assembly was of slightly higher completeness than Murphy (Table 1), so we designate Maya as the *I. ricinus* reference genome assembly.

The genome assemblies of *I*. *ricinus* Maya and Murphy had a similar number of gene predictions, with approximately 27,000 genes and 204,000 exons (Table 1B). While the number of genes for *I*. *ricinus* Maya and Murphy is less than the *I. scapularis* genome assembly (Table 1B), most likely this is due to the *I. scapularis* genome being annotated with NCBI’s RefSeq pipeline rather than a biological difference. The gene density per Gbp in *I*. *ricinus* is approximately 12,600 and there is an average of 7.5 exons per gene. The average intron length for *I*. *ricinus* Maya was 3,308 bp and for *I*. *ricinus* Murphy 3,296 bp. The total coding sequence length for *I. ricinus* Maya is 37,691,999 bp and *I. ricinus* Murphy 37,663,835 bp, which is only 1.76 % of both genomes. In addition, the introns are 24.63% of the *I. ricinus* Maya genome and 24.53% *I. ricinus* Murphy genome.

### Comparative genome analysis of key Lyme borreliosis tick vectors (*Ixodes* species)

We compared the whole genome assemblies of the tick vector for Lyme borreliosis in Western Eurasia (*I. ricinus*) and North America (*I. scapularis*). Direct mapping of the assembled *I. ricinus* Maya and Murphy against *I. scapularis* gave an average identity of 84%.

As a chromosome-anchored *Ixodes* genome is not yet available, we mapped the *I*. *ricinus* contigs against the 14 chromosome-scale scaffolds of *I*. *scapularis* and each *I*. *ricinus* scaffold was labelled with a unique letter (Figure 1 and Figure S2-S4). A total of 18.6% (53% of contigs) and 17.3% (54% of contigs) of the total genome length were not included in the chromosome-scale scaffolds from *I*. *ricinus* Maya and Murphy, respectively.

**Figure 1.**
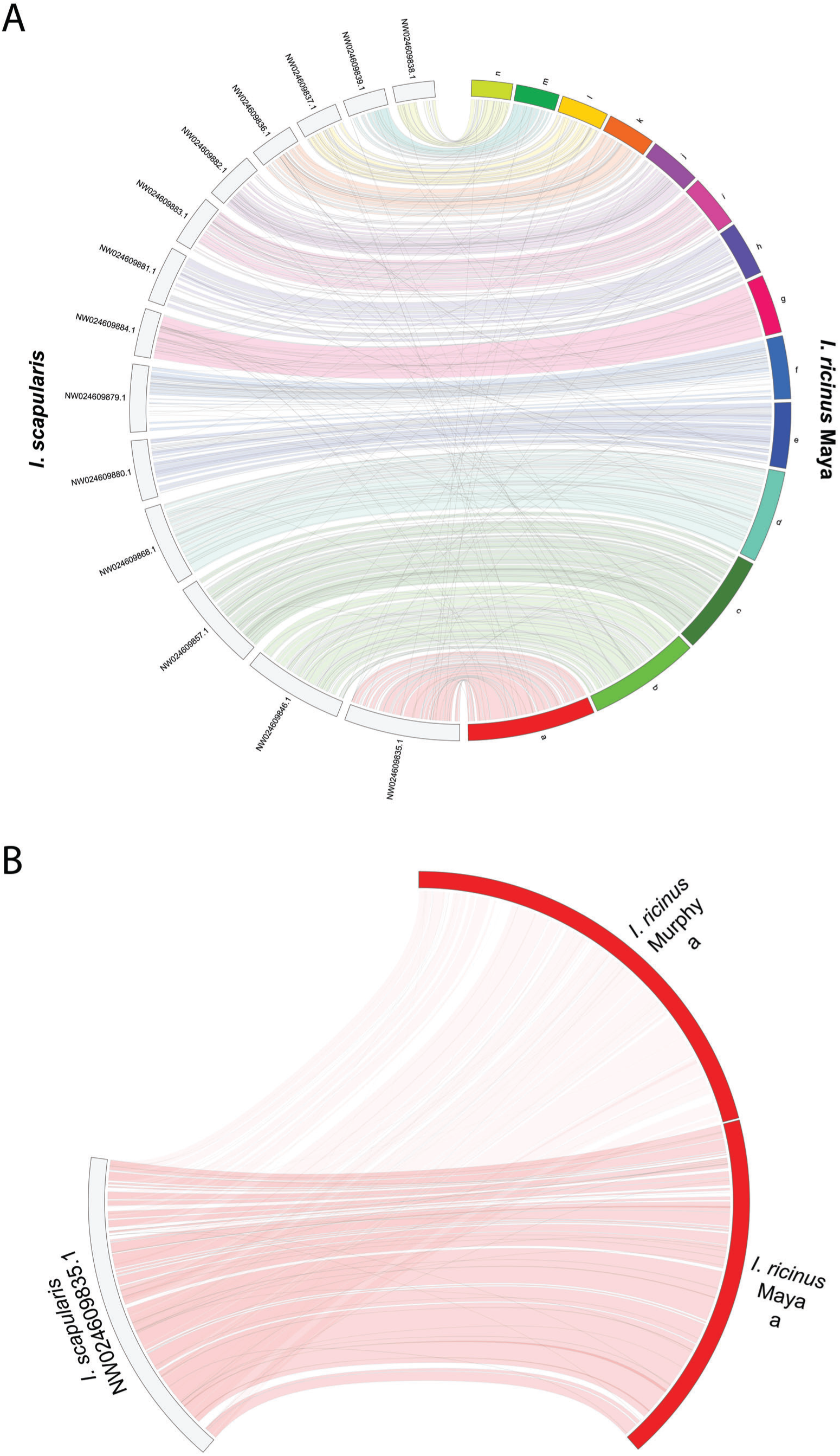
Chromosome-scale scaffold synteny of *Ixodes scapularis* and *I*. *ricinus*. (A) The 14 chromosome-scale scaffolds of *I. scapularis* (gray) with the 14 corresponding scaffolds of *I. ricinus* Maya (colored); (B) the largest scaffold of *I. scapularis* with scaffold “a” of *I. ricinus* Maya and *I. ricinus* Murphy. Gray lines denote the repetitive hits.

The two *I. ricinus* genome assemblies and the *I. scapularis* genome share 13,963 orthogroups (Figure S5). These shared orthogroups are determined by 18,468 transcripts in *I. ricinus* Maya, 18,487 transcripts in *I. ricinus* Murphy and 22,206 transcripts in *I. scapularis*. There are 14,942 unique gene IDs in *I*. *ricinus* Maya for the shared orthogroups, 14,918 in *I*. *ricinus* Murphy and 17,807 in *I. scapularis*. Within the shared orthogroups there are 9,703 that have a single transcript for all three genomes, all other shared orthogroups have varying transcript amounts among the species. Looking at orthogroups unique to each *Ixodes* species, there are 4,408 *I. ricinus*-specific orthogroups which are made up of 4,619 transcripts for Maya with 4,119 unique gene IDs associated with them, whereas Murphy consists of 4,588 transcripts with 4,084 unique gene IDs. There are 856 unique orthogroups to *I. scapularis* made up of 3,340 transcripts, and all orthogroups were determined by multiple transcripts with 2,460 unique gene IDs. While the orthogroups unique to *I. ricinus* genomes had 1,008 orthogroups that were determined by a single transcript each.

### Annotation of repetitive DNA in *Ixodes* ticks and creation of a tick repeat library

Currently an “*Ixodes*” query in RepBase, the key repository of TE consensus sequences, retrieves only 65 entries and 77% of these entries belong to a single Long Terminal Repeat (LTR) superfamily, Gypsy. Using RepeatModeler we recovered over 4,500 putative TE consensus sequences across the two *I. ricinus* assemblies and the *I. scapularis* assembly. Our final tick repeat library contains 3,684 consensus sequences that met the 80-80-80 rule^28^ and were therefore classified as belonging to families assignable to known TE orders and families (Table S1).

The *Ixodes* repeat library has 30 TE superfamilies (Table S1). The library is dominated by Terminal Inverted Repeat (TIR) transposons (43% of the library). The most numerous superfamily in the library was TIR-like, which have consensus sequences that harbor TIRs but do not have any obvious target site duplications (29% of the entire library and 67% of TIRs in the library). Gypsy LTR elements were the second most prevalent superfamily in the library (15% of the entire library) and the third was R1 Long INterspersed Elements (LINEs) (8% of the entire library).

### Characterization of the repetitive *Ixodes* genome

We find the *I*. *ricinus* genomes are at least 73% repetitive DNA, of which 69% is TEs (Figure 2 and Figure S6A, Table S1). The *I*. *scapularis* genome is at least 74% repetitive DNA, of which 68% is TEs (Figure S6B, Table S1). Our genome proportions for repetitive DNA in *Ixodes* are slightly higher than identified in previous studies^20,23–25,27^, likely due to their use of traditional annotation pipelines, which use homology-based approaches or apply the raw output of TE-finders to the genome assembly. These traditional pipelines underestimate TE content, especially for non-model species^29,30^.

**Figure 2.**
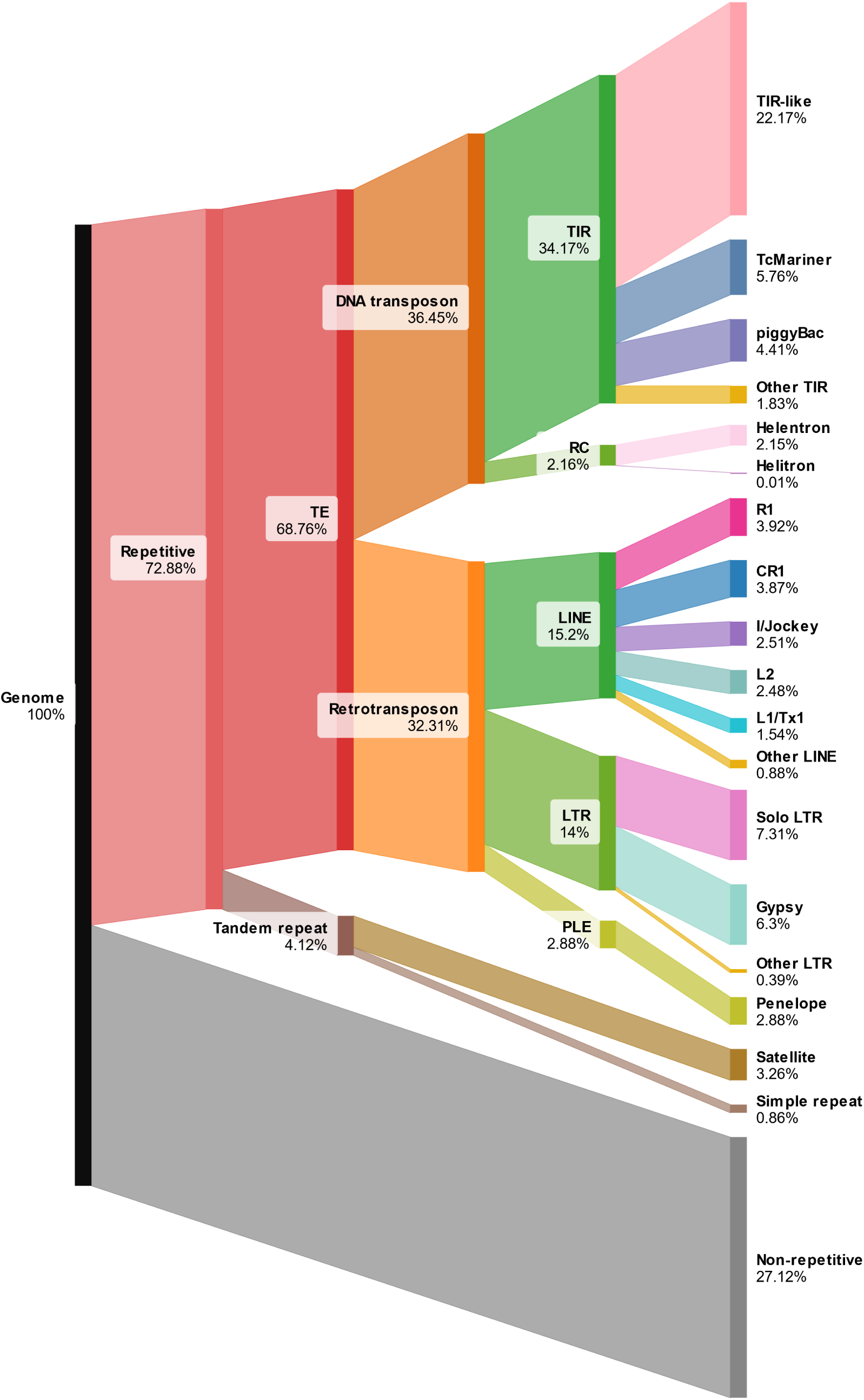
Genome proportions of repetitive and non-repetitive DNA in *Ixodes ricinus* Maya. The majority of the *I. ricinus* genome is repetitive DNA, dominated by transposable elements (TEs): Terminal Inverted Repeat (TIR) transposons; Long INterspersed Element (LINE) retrotransposons; Long Terminal Repeat (LTR) retrotransposons; Penelope-like (PLE) retrotransposons; and rolling-circle (RC) transposons. Genome proportions for *I. ricinus* Murphy and *I. scapularis* see supplemental material Figure S6.

The TEs in the *Ixodes* genome assemblies (Table S1) are dominated by TIR transposons (34-35% of each genome), predominantly the aforementioned TIR-like elements (21-22%), but also the TcMariner and piggyBac superfamilies. The next most common TEs are LINE retrotransposons (15% of each genome), mostly CR1, R1, I/Jockey, L2, and L1/Tx1 superfamilies. The third most common TEs are LTR retrotransposons (13-14% of each genome), which are predominantly solo LTRs and members of the Gypsy superfamilies. Fourth are Penelope-like Element (PLE) retrotransposons (3% of each genome). The fifth most common TEs are the rolling-circle (RC) transposons (2% of each genome), mainly Helentrons.

The tandem repeats in the *Ixodes* genome assemblies (Table S1) are dominated by satellite DNA (3-4% of each genome). This satellite DNA is often associated with non-LTR LINEs, comprising approximately one quarter of the total satellite content in *I*. *ricinus* and one third in *I. scapularis*.

### Species specific accumulation patterns of TEs in *Ixodes*

Recent TE expansions in the *Ixodes* genomes have occurred, indicated by the left-skew of the TE landscapes (Figure 3). The TE accumulations in the two *I. ricinus* genome assemblies are an almost exact match, with only minor differences in the genome proportions (Figure 3). In contrast, post-divergence of *I. ricinus* and *I. scapularis* each species has a distinct TE accumulation landscape. For example, most recently in *I. ricinus* there has been an increase in Helentron elements, whereas this superfamily has lower rates of accumulation in *I. scapularis* (Figure 3). Also, most recently in *I. ricinus* there has been a decrease in TcMariner accumulation, which is not seen in *I. scapularis* (Figure 3). In *I. ricinus* the Sola superfamily has been largely dormant throughout its evolutionary history and in *I. scapularis* this superfamily recently expanded (Figure 3).

**Figure 3.**
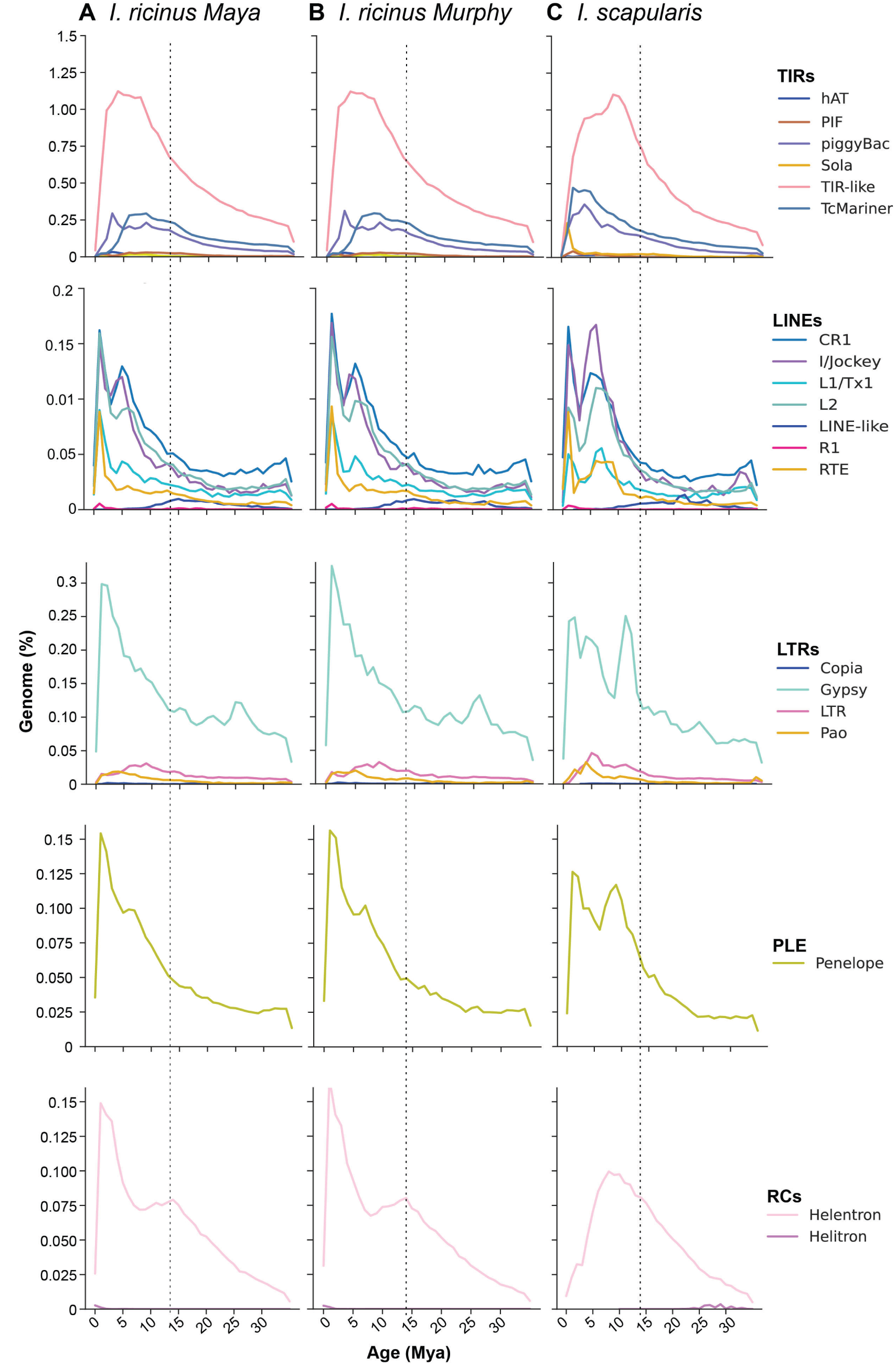
Transposable element timeline for *Ixodes* species. (A) *Ixodes ricinus* Maya, (B) *I. ricinus* Murphy, and (C) *I*. *scapularis* reference genome. The genome percentage of each TE superfamily was calculated using the total TE superfamily content in bp for each age bin and dividing by the total genome size. The estimated time of accumulation is based on a neutral mutation rate of 7.68 x 10^-9^ substitutions/site/million years and divergences between individual elements and their respective consensus sequences. Each row designates a TE order (TIR DNA transposons (TIRs); non-Long Terminal Repeats retrotransposons (non-LTRs); Long Terminal Repeats retrotransposons (LTRs); Penelope-like elements retrotransposons (PLEs); and rolling circle transposons (RCs). The vertical dashed line indicates the approximate divergence time between *I. ricinus* and *I. scapularis* of 13 million years ago^31^.

## DISCUSSION

We have generated two high-quality genome assemblies for the major disease vector *I*. *ricinus*. These assemblies have comparable quality to the genomes of other tick species such as *I*. *scapularis*^20,21^ and *I*. *persulcatus*^24^. The genome size of *I*. *ricinus* is approximately 2.15 Gbp. This size is many fold larger than the genomes of the most well-studied insects *Drosophila melanogaster* 0.14 Gbp^32^ and *Anopheles gambiae* 0.26 Gbp^33^, plus larger than closer related arthropods such as *Varroa destructor* 0.37 Gbp^34^ and *Parasteatoda tepidariorum* 1.4 Gbp^35^. The large genome size of *Ixodes* ticks is largely due to the high proportion of repetitive DNA their genomes contain, which is commonly found in arthropods with gigantic genome sizes^22,36–38^.

Our repetitive DNA curation finds nearly three quarters of the *I*. *ricinus* genome assemblies are repeat-derived, with almost all the repetitive DNA being derived from TEs. *Ixodes* ticks have one of the highest proportions of TEs currently reported for an animal genome^22,29,38^. Other tick species also have high proportions of TEs, ranging from over 25% to 64% of the genome^24,39^. In contrast, the proportion of TEs in mosquitoes can range from almost zero to 50% of the genome^40^.

TE-driven genome evolution^41^ has occurred in *Ixodes* ticks relatively recently, within the last ∼13 million years^31^. In two sister lineages, *I. ricinus* and *I. scapularis*, we find the TE proportions have distinctive post-speciation trajectories. The TEs in these tick species are often highly active and rapidly turned over, suggesting that TEs accumulate and then are quickly removed from the genomes via nonhomologous recombination^37,42–45^. These phenomena suggest rapid structural change in the genomes once the two *Ixodes* species separated.

TEs are known drivers of species diversity^41,46–52^ and we propose this is occurring for *Ixodes* ticks. Notably, the tick genus with the largest proportion of TEs in its genome, *Ixodes*, is also the most speciose tick genus^53^, connecting TEs as potential drivers of genome and species diversification^54,55^. However, there is typically a positive relationship between TE content and genome size^56^. Given this relationship, one would expect the other tick genera that have larger genomes than *Ixodes*^24^ to exhibit higher proportions of TEs. As our analysis of *Ixodes* TEs is the most extensive and detailed examination of tick TEs to date, previous analyses may have underestimated TE content in other tick species.

The abundance and diversity of TEs in the genomes of ticks could be due to their obligate blood-feeding parasitic lifestyle. Ticks have a close association with their vertebrate hosts (from which they take blood at least three times throughout their lifecycle) and subsequently contain a plethora of microbiota. These close associations are reflected in our *I*. *ricinus* sequencing reads, as we found genome contamination from the tick’s vertebrate host and microbiota. We suggest tick genomes, with their close associations to the DNA of other organisms, may be a key horizontal transfer point for TEs. There is evidence that horizontal transfer of TEs can occur from eukaryote parasites to their hosts and viruses to eukaryotes^57–61^. For example the LINE BovB was likely horizontally transferred to different vertebrates via a tick^62^. Also, the blood-feeding behavior of a parasite might be important for TE abundance and diversity, as not all blood-feeders have high proportions of TEs^40^. We suggest future studies should investigate whether tick species and blood-feeding parasites that are host-generalists have greater TE diversity compared to those that are host-specialists.

Our genome assemblies for the tick *I*. *ricinus* will facilitate a deeper understanding of the fundamental biology of *Ixodes* species in general and *I*. *ricinus* specifically. Comparative genomics for ticks will be further enhanced with chromosome-scale scaffolding for *I*. *ricinus*^63^ and population genetic studies^c.f.64^. More broadly our findings indicate further studies of TEs in ticks and blood-feeding parasites are required for a more comprehensive understanding of the evolutionary history of these medically and veterinary important taxa.

## MATERIALS AND METHODS

### Tick samples

We collected questing *I*. *ricinus* nymphs in September 2020 by blanket dragging in De Dorschkamp, a deciduous forest between Wageningen and Renkum, the Netherlands (51°58’39.5“N, 5°41’59.3“E). These nymphs were morphologically identified to species level using an identification key^65^. To obtain adult *I*. *ricinus* we fed the nymphs on an artificial membrane blood feeding system, as described in ^66^. The bovine blood was obtained from Carus (Wageningen University, The Netherlands), under animal ethics protocol no. AVD1040020173624. Engorged nymphs were collected from the feeding membrane and incubated at room temperature until they molted to adults. The adults were collected and stored at -80° C. We designated *I*. *ricinus* adult female #1 as biospecimen Maya1009 and *I*. *ricinus* adult female #2 as biospecimen Murphy0812.

### DNA isolation and genome sequencing (Oxford Nanopore Technologies and Illumina)

DNA extraction and sequencing was performed at Future Genomics Technologies BV (Leiden, the Netherlands). The two individual, whole *I. ricinus* adults were ground to a fine powder using a pestle and mortar with liquid nitrogen. To extract high molecular weight DNA a Genomic-tip 20/G kit (Qiagen Benelux BV, Venlo, the Netherlands) was used according to the standard protocol. DNA quality was measured via electrophoresis in Genomic DNA ScreenTape on an Agilent 4200 TapeStation System (Agilent Technologies Netherlands BV, Amstelveen, the Netherlands) and total DNA measured using a Qubit 3.0 Fluorometer (Life Technologies Europe BV, Bleiswijk, the Netherlands); we obtained between 3.05 μg to 5.96 μg of total DNA.

The DNA samples were used to prepare a 1D ligation library using the Ligation Sequencing Kit SQK-LSK110 according to the manufacturer’s instructions (Oxford Nanopore Technologies, Oxford, United Kingdom). Oxford Nanopore Technologies (ONT) libraries were first tested on a MinION flowcell (FLO-MIN106) and subsequently run on an R9.4.1 PromethION flowcell (FLO-PRO002) using the following settings: basecall model: high-accuracy; basecaller version: 4.0.11 (PromethION).

Parallel aliquots of the DNA samples used for ONT sequencing, were used to prepare Illumina libraries using the Nextera DNA Flex Library Prep Kit according to the manufacturers’ instructions (Illumina Inc. San Diego, CA, USA). Library quality was measured via electrophoresis in D1000 ScreenTape on an Agilent 4200 TapeStation System (Agilent Technologies Netherlands BV, Amstelveen, the Netherlands). The genomic paired-end (PE) libraries were sequenced with a read length of 2 x 150 nt using the Illumina NovaSeq 6000 system. The ONT sequences underwent quality assessment using Nanoplot v1.42^67^ and Illumina sequences were assessed using MultiQC v1.14^68^ .

### Genome assembly and quality assessment

The genome sequence data of Maya1009 and Murphy0812 were submitted to a custom hybrid genome sequence pipeline (Figure S7). First, ONT reads were filtered using yacrd v.0.6.2^69^ to remove potential chimeric reads generated by the sequencing ligation kit. Then bovine blood meal genome contamination was removed from the filtered reads using a custom database containing *Bos taurus* (GCA_002263795.3). The ONT reads were then assembled using Flye 2.9.1^70^. The final scaffolds generated by the pipeline was then polished using Illumina short reads and NextPolish v1.4.0^71^ with three reruns. To predict the putative haploid genome size of the assemblies, we used purge_haplotigs v.1.1.2^72^, to remove redundant contigs.

To check for any genome contamination on the polished deduplicated assemblies we used NCBI’s Foreign Contamination Screen tool^73^, gx-db version 0.4.0 and tax-id 6944 for *Ixodes* species. This tool checks if contamination is detected as a contig or within an assembled contig. Any contaminant we detected in the assemblies was removed or hardmasked using bedtools v2.30.0^74^. To quantify the amount of biological contamination we calculated the proportion of blood meal and microbiota reads in the two genome assemblies.

To assess the completeness of the two assembled genomes, we submitted them to Benchmarking Universal Single-copy Orthologs (BUSCO) v5^75^ using the Arthropoda database. In parallel, the genomes were annotated using a combination of AUGUSTUS v3.4.0^76^ gene prediction and Liftoff v1.6.3^77^ annotation transfer tool using the recent *I*. *scapularis* genome assembly annotation (NCBI Assembly GCA_016920785.2^20^) as a template.

### Gene and gene family annotation

The annotated Maya1009 and Murphy0812 gff files, plus the gff file associated with the current *I. scapularis* assembly (NCBI Assembly GCA_016920785.2^20^) were submitted to agat_sp_extract_sequences.pl from the AGAT v.1.0 pipeline^78^ to obtain all annotated protein sequences in a FASTA. Those protein FASTA files were used to run an ortholog comparison for the *I. ricinus* samples (Maya1009, Murphy0812) using Orthofinder v.2.5.2^79^. The overlap of the resulting Orthogroups were then plotted using R package VennDiagram v.1.7.3^80^. To examine the gene families that were unique to the *Ixodes* species, we ran Orthofinder v.2.5.2 to compare the current *I. scapularis* genome and the *I. ricinus* reference assembly (Maya1009). Genes that were only found in the *I. ricinus* sample Maya1009 were extracted from the dataset with the corresponding transcript information. These steps were repeated for the orthogroups unique to the *I. scapularis* genome.

### Whole genome comparison

We compared the Maya1009 and Murphy0812 genome assemblies to the current reference genome assembly of *I. scapularis* (NCBI Assembly GCA_016920785.2^20^), which is the most closely related tick species with a genome assembly and chromosome-scale scaffolds available. The average identity between the *I. ricinus* genomes and *I. scapularis* was calculated using FastANI v1.33^81^. To obtain an overall comparison at a chromosome-scale level between the two *Ixodes* species, we first hard masked for repeats in both *I. ricinus* genomes using RepeatModeler v2.0.3^82^. Then the masked *I*. *ricinus* contigs were “scaffolded” to the 14 chromosome-scale scaffolds (i.e. the 14 longest scaffolds) from the *I. scapularis* reference genome (representing almost 90% of the *I. scapularis* total genome size^20^) using RagTag v.2.1.0^83^. This step was required because the majority of the *I. ricinus* scaffolds are shorter than the *I. scapularis* reference genome scaffolds. The scaffolded contig IDs are available in Table S2A and B for Maya1009 and Murphy0812, respectively. We conducted synteny analysis using minimap2 v.2.2.24^84^ and Circos^85^ was used to make the synteny plots.

### Repetitive DNA identification and characterization

The repetitive DNA in *Ixodes* ticks was subjected to detailed manual annotation. We had previously characterized the repetitive DNA of *I. scapularis* using a standard RepeatModeler/RepeatMasker pipeline^20^. Here we generated a *de novo* repeat library, using a custom pipeline, for the genome assemblies of *I*. *ricinus* (Maya1009 and Murphy0812) and *I. scapularis*^20^. First we used RepeatModeler v.2.0.3^82^ to produce putative consensus sequences that we extracted (https://github.com/davidaray/bioinfo_tools/blob/master/extract_align.py) and subjected to extension using RAM^86^. The putative consensus sequences were categorized as a LTR retrotransposon, LINE retrotransposon, Short INterspersed Element (SINE) retrotransposon, PLE retrotransposon, TIR transposon, RC transposon, Maverick transposon or unidentified. This categorization was done using a custom bash script (TEcurate.sh) of RepeatClassifier (part of the RepeatModeler package) and BLASTP searches (e-values >1e^-50^) against a database of known autonomous TEs. We then used a custom curation script (https://github.com/davidaray/bioinfo_tools/blob/master/TEcurate.sh) and the TE-Aid (https://github.com/clemgoub/TE-Aid) package to generate genome coverage plots, self-alignment dot-plots, structure and ORF plots, and copy number estimates. For sequences categorized as ‘unidentified’, we determined likely group membership using the TE-Aid plots to identify structural hallmarks (i.e. TIRs and LTRs) and sequence characteristics (repetitive tails, Helitron-specific CTRR motifs, SINE A-B boxes, piggyBac TTAA target site duplications, etc.), if the TE remained unidentified it was categorized as unknown. The resulting tick repeat library was combined with known arthropod TEs in RepBase and the TEs we had previously identified in *I. scapularis*^20^ to create a custom library that was applied to Maya, Murphy, and the *I. scapularis* assembly. This combined library was subjected to analysis using cd-hit-est and a 90% similarity cutoff to eliminate highly similar consensus sequences.

For the TEs we used the identifiers provided by TEcurate.sh to generate a unique identifier that included the species of origin, RepeatModeler ID, and TE Class/Family information. For example, iRic1.5.2409#LINE/I: identified in Maya1009 (Murphy0812 is designated as iRic2), RepeatModeler ID rnd-5_family-2409 (5.2409) and RepeatClassifier/blastp analysis identified it as a LINE element of the I family. For the unidentified TEs we used visual examination and categorization to assign an appropriate identifier.

We determined the TE consensus sequences. The LTR and PLE retrotransposons we identified were subjected to postprocessing to obtain consensus sequences. LTRs were processed by hand to subdivide them into their LTR and internal segments^87^. PLEs often insert as tandemly repeated arrays, so we used the TE-Aid output to split the tandem arrays into a single representative full-length consensus. RAM occasionally falsely identifies segmental duplications as TEs. Therefore, any low-copy (<50 copies) consensus sequences greater than 10 kb and with no consistent TE structural hallmarks were assumed to be segmental duplications or assembly artifacts and removed from the library. Consensus sequences with fewer than 10 full-length copies were also removed from the library. We submitted all the novel TE consensus sequences we identified to the Dfam TE database.

We identified that the LINE superfamilies CR1 and L2 were commonly associated with satellite repeats at their 5’ ends. Ten randomly selected consensus sequences were subjected to further characterization to check whether these repeats were an artifact of genome assembly. The ONT reads were searched using 400 bp queries consisting of 200 bp of the upstream satellite sequence and 200 bp of the downstream LINE sequence. In all ten test cases, the queries were identified in the original reads, confirming that these satellite/LINE chimeras exist in the genome of these ticks. To aid with submission to the Dfam repeat database, we separated the satellite sequences from the LINE consensus sequences and included both in the final tick repeat library. For example, the CR1 consensus iRic2.5.1902 was subdivided into its satellite component (iRic2.5.1902S) and its LINE component (iRic2.5.1902L). These satellite repeats are a significant portion of the consensus sequences. For example, in iRic2.5.1902 the satellite consists of a 316 bp repeated 13.2 times and comprises more than half of the original consensus (4,211 bp versus 3,760 bp derived from the CR1 portion).

Our tick repeat library was used to mask the *I. ricinus* and *I. scapularis* genome assemblies with RepeatMasker. The output was processed to eliminate any overlapping hits using RM2Bed.py, part of the RepeatMasker installation package, generating BED files for downstream analyses. In addition, we used the tick repeat library to calculate the proportion of each TE superfamily in the three genomes and then created a Sankey diagram using SankeyMATIC.

We obtained TE accumulation time estimates by estimating a neutral mutation rate for *Ixodes*. We aligned randomly selected orthologous introns (see Supplemental Information Item 1) from *I. ricinus* and *I. scapularis* in MEGA v11^88^ under the maximum composite likelihood model (0.999). Then, using a species divergence time of 13 million years ago^31^, we calculated a neutral mutation rate of ∼7.68 x 10^-9^ substitutions/site/million years for *Ixodes*.

## ETHICS APPROVAL STATEMENT

The bovine blood, which was used for an artificial membrane blood feeding system was obtained from Carus (Wageningen University, The Netherlands), and the procedures were approved by the Animal Ethics Committees of Wageningen Research under animal ethics protocol no. AVD1040020173624. The animal experiments were approved by the Dutch Central Authority for Scientific Procedures on Animals.

## AVAILABILITY OF DATA AND MATERIALS

Raw reads and the genome assemblies are available under the BioProject PRJNA816462 and Biosamples (Maya1009/SAMN26676983 and Murphy0812/SAMN26677070).

Tick consensus sequences and supporting information have been deposited in the Dfam transposable element database (https://www.dfam.org/).

## ACKNOWLEDGEMENTS

We thank Ron Dirks (Future Genomics Technologies, Leiden, The Netherlands) for invaluable advice on sample preparation and next generation sequencing.

## AUTHOR CONTRIBUTIONS

Conceptualization: IR, HS, TCG

Validation: RPB, NSP, JCF, KCD, DAR

Formal analysis: RPB, NSP, JCF, KCD, DAR

Investigation: TA, JWB

Resources: TA, JWB

Data Curation: RPB, NSP, DAR

Writing - Original Draft: IR, DAR

Writing - Review & Editing: All

Visualization: IR, RPB, NSP, JCF, DAR

Supervision: HS, DAR, TCG

Project administration: IR, RPB, HS, DAR, TCG

Funding acquisition: HS

