## Supplementary material for "The repetitive genome of the *Ixodes ricinus* tick reveals transposable elements have driven genome evolution in ticks": Figure S1; Figure S2; Figure S3; Figure S4; Figure S5; Figure S6; Figure S7

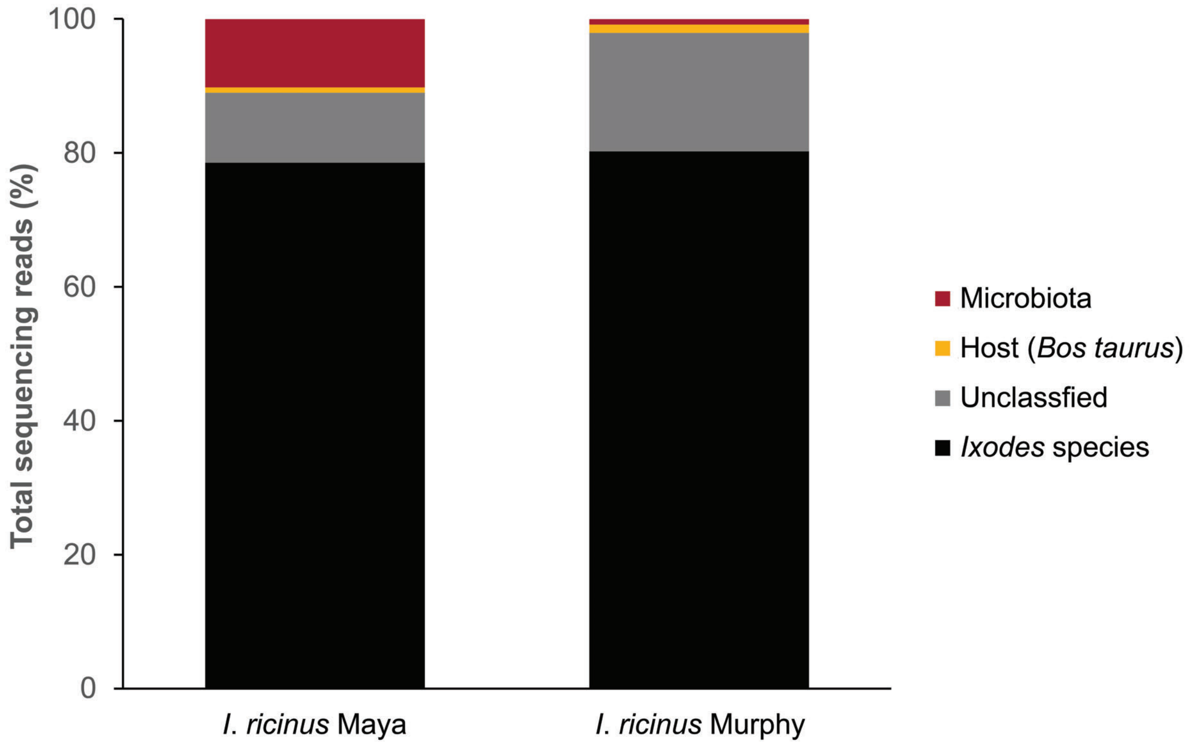


**Figure S1. Taxonomic classification of sequencing reads from *Ixodes ricinus* Maya and Murphy to identify biological contamination.** We retained reads classified as *Ixodes* species or reads that remained unclassified, as they were considered to be potential *I. ricinus* genomic sequences. Filtered from the genome assemblies was any biological contamination, either reads classified as host (*Bos taurus*) or the reads from contigs classified as microbiota.

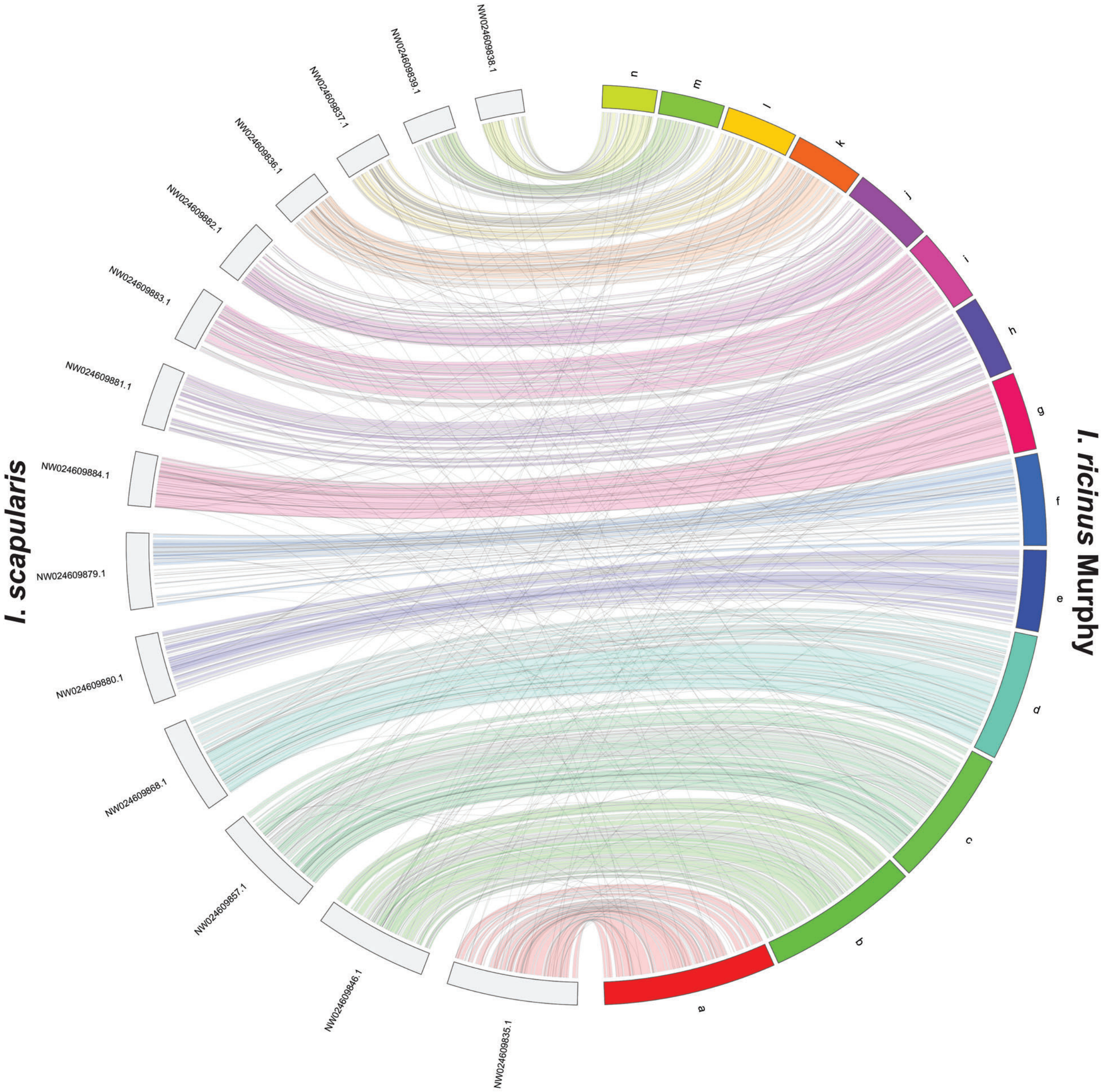


**Figure S2. Chromosome-scale scaffold synteny of *Ixodes scapularis* and *I. ricinus* Murphy.** The 14 chromosome-scale scaffolds of *I. scapularis* (gray) with the 14 corresponding scaffolds of *I. ricinus* Murphy (colored). Gray lines denote the repetitive hits.

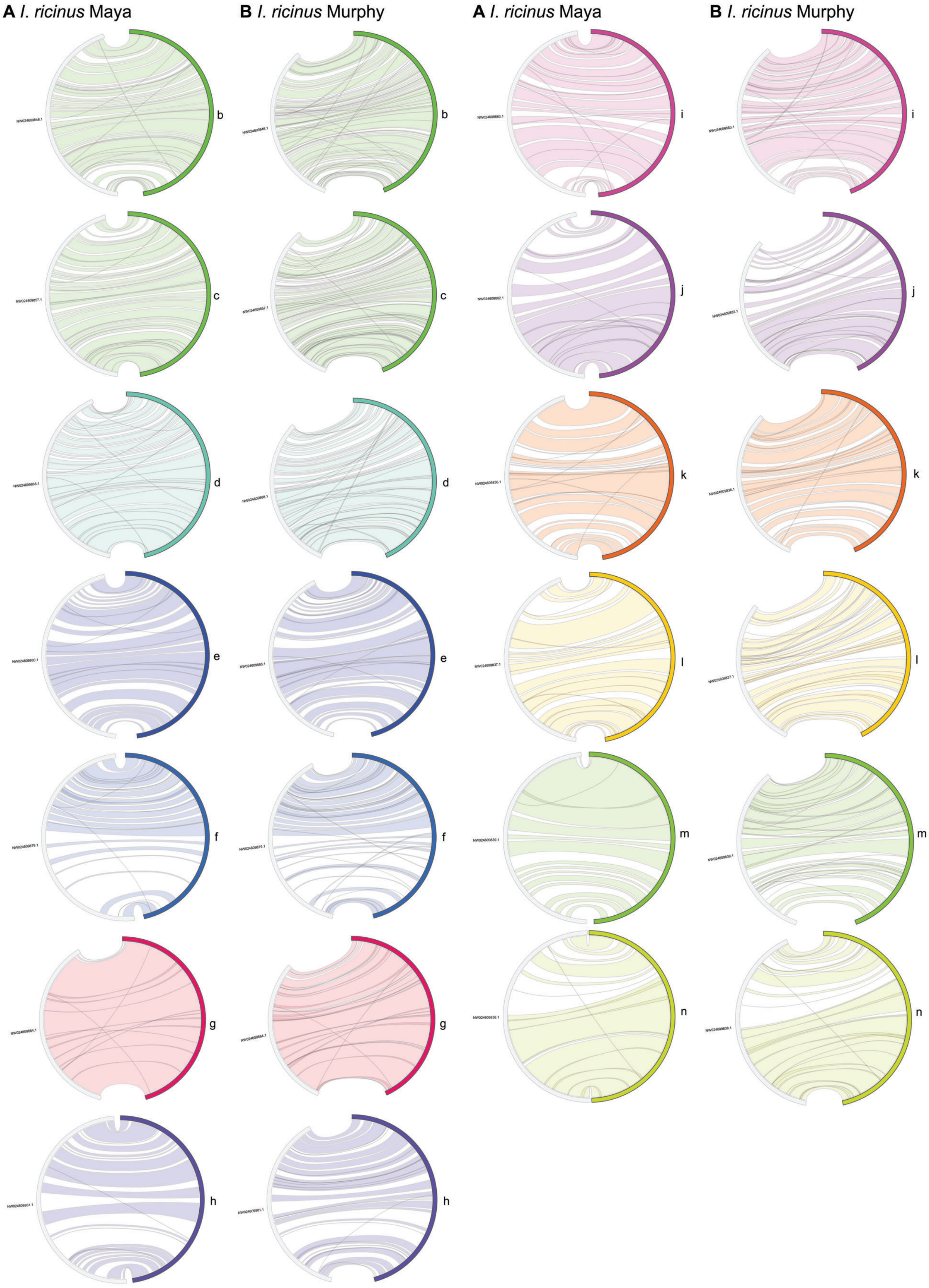
A I. ricinus Maya
B I. ricinus Murphy
A I. ricinus Maya
B I. ricinus Murphy
b
c
d
e
f
g
h
i
j
k
l
m
n

**Figure S3. Chromosome-scale scaffold synteny of *Ixodes scapularis* with the two *I. ricinus* genome assemblies.** The second to fourteenth largest scaffolds of *I. scapularis* (the largest is in Figure 1B) with the corresponding scaffolds of (A) *I. ricinus* Maya and (B) *I. ricinus* Murphy, the letters “b” to “n” refer to the *I. ricinus* scaffolds. Gray lines denote the repetitive hits.

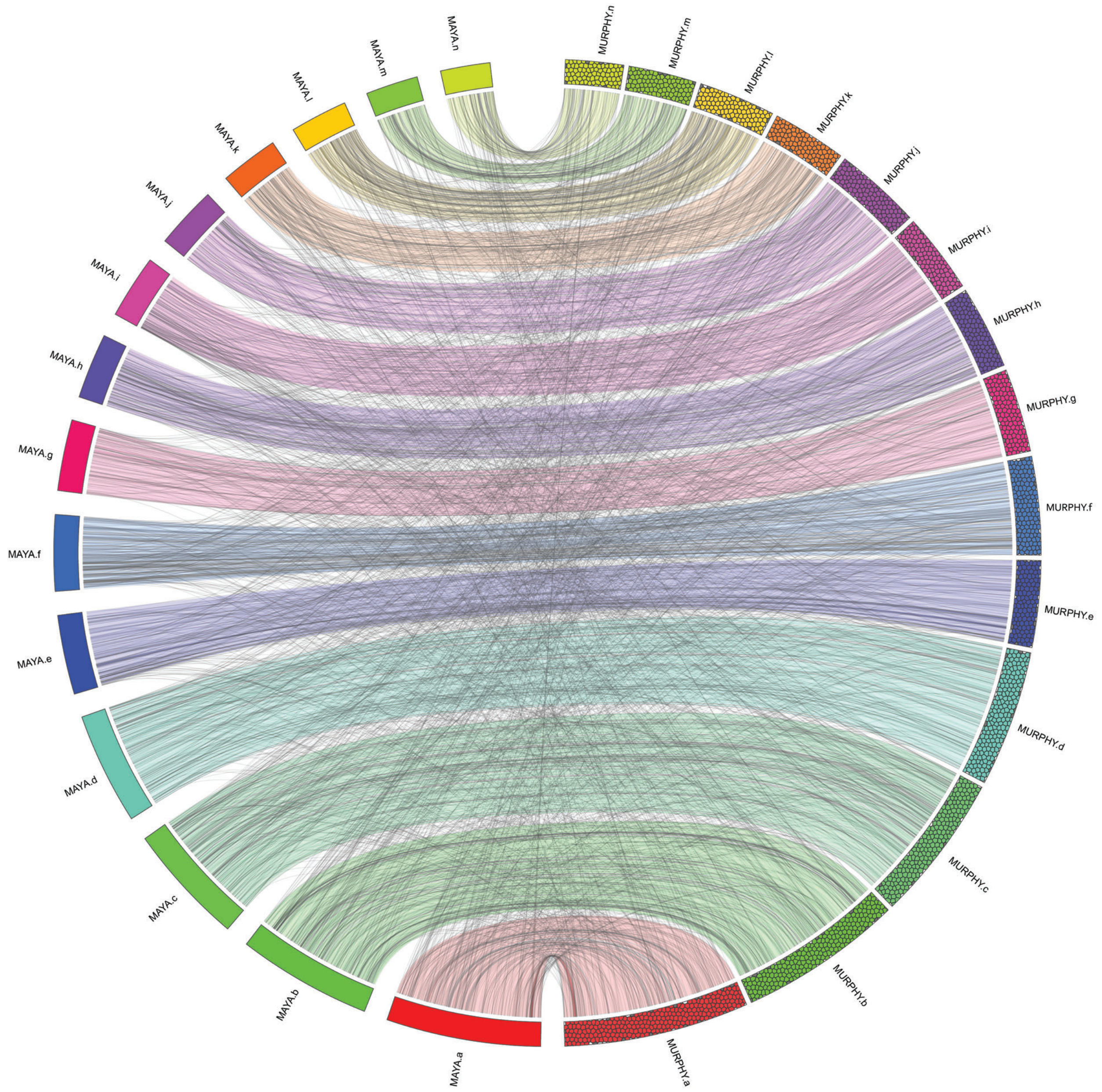


**Figure S4. Chromosome-scale scaffold synteny of the two *Ixodes ricinus* genome assemblies.** The 14 chromosome-scale scaffolds of *I. ricinus* Maya (colored) with the 14 corresponding scaffolds of *I. ricinus* Murphy (colored bubbles). Gray lines denote the repetitive hits.

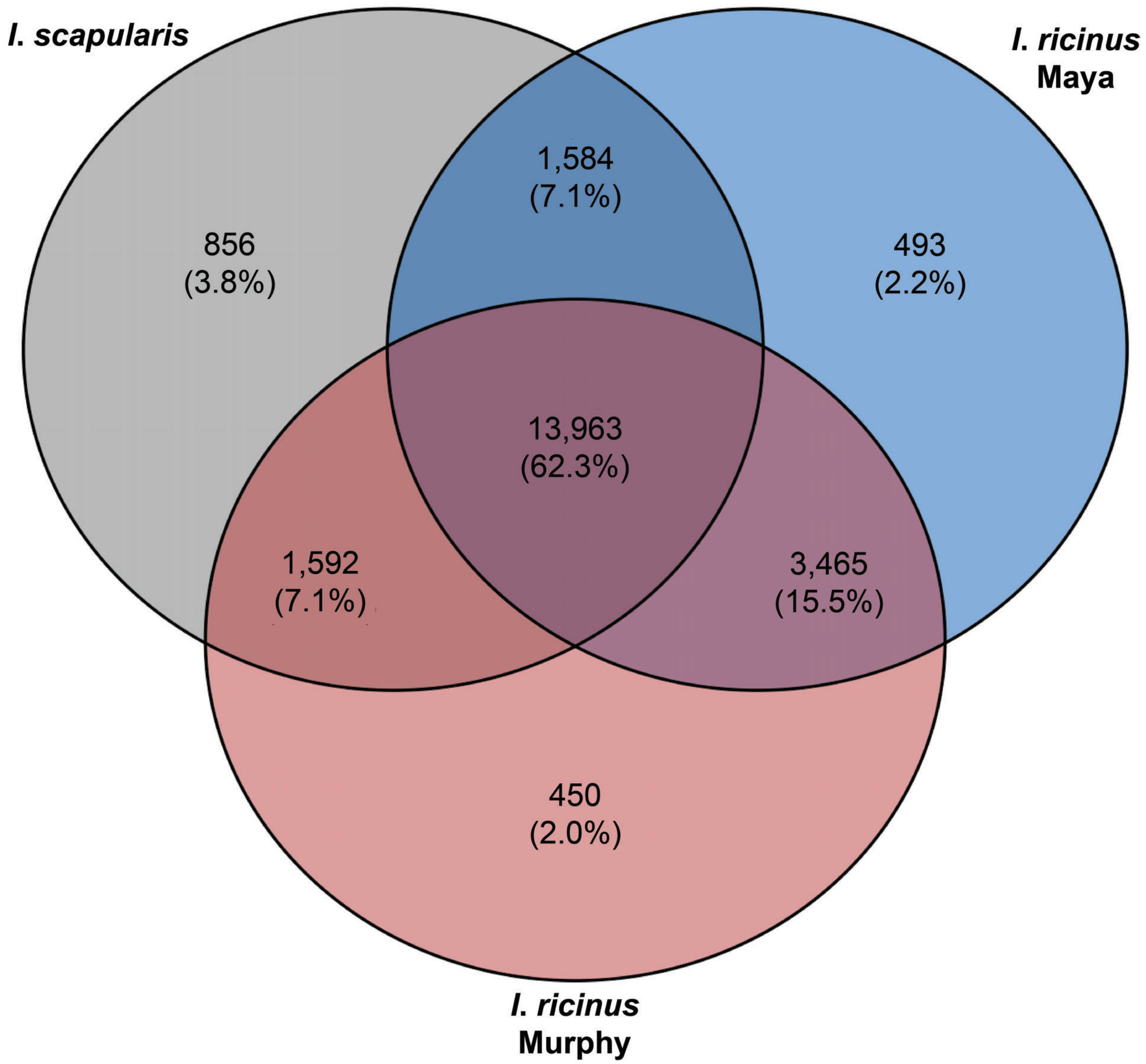


**Figure S5. Gene orthology in *Ixodes* species.** Orthogroups for *Ixodes ricinus* Maya (blue), *I. ricinus* Murphy (red) and the *I. scapularis* reference genome (gray), highlighting shared and unique orthogroups.

37 **A**

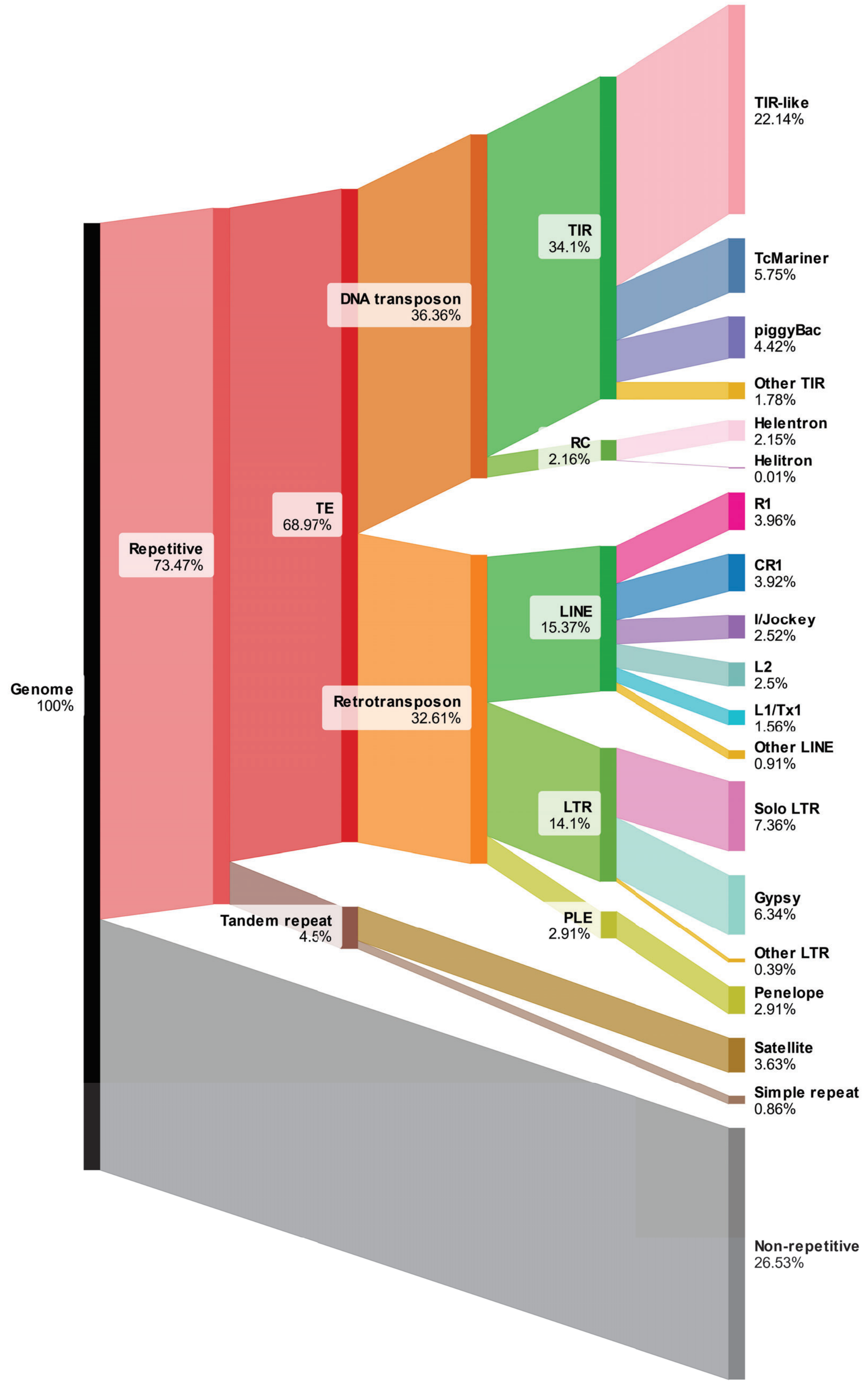

Genome
100%
Repetitive
73.47%
TE
68.97%
DNA transposon
36.36%
Retrotransposon
32.61%
Tandem repeat
4.5%
TIR
34.1%
RC
2.16%
LINE
15.37%
LTR
14.1%
PLE
2.91%
TIR-like
22.14%
TcMariner
5.75%
piggyBac
4.42%
Other TIR
1.78%
Helentron
2.15%
Helitron
0.01%
R1
3.96%
CR1
3.92%
I/Jockey
2.52%
L2
2.5%
L1/Tx1
1.56%
Other LINE
0.91%
Solo LTR
7.36%
Gypsy
6.34%
Other LTR
0.39%
Penelope
2.91%
Satellite
3.63%
Simple repeat
0.86%
Non-repetitive
26.53%


38

B

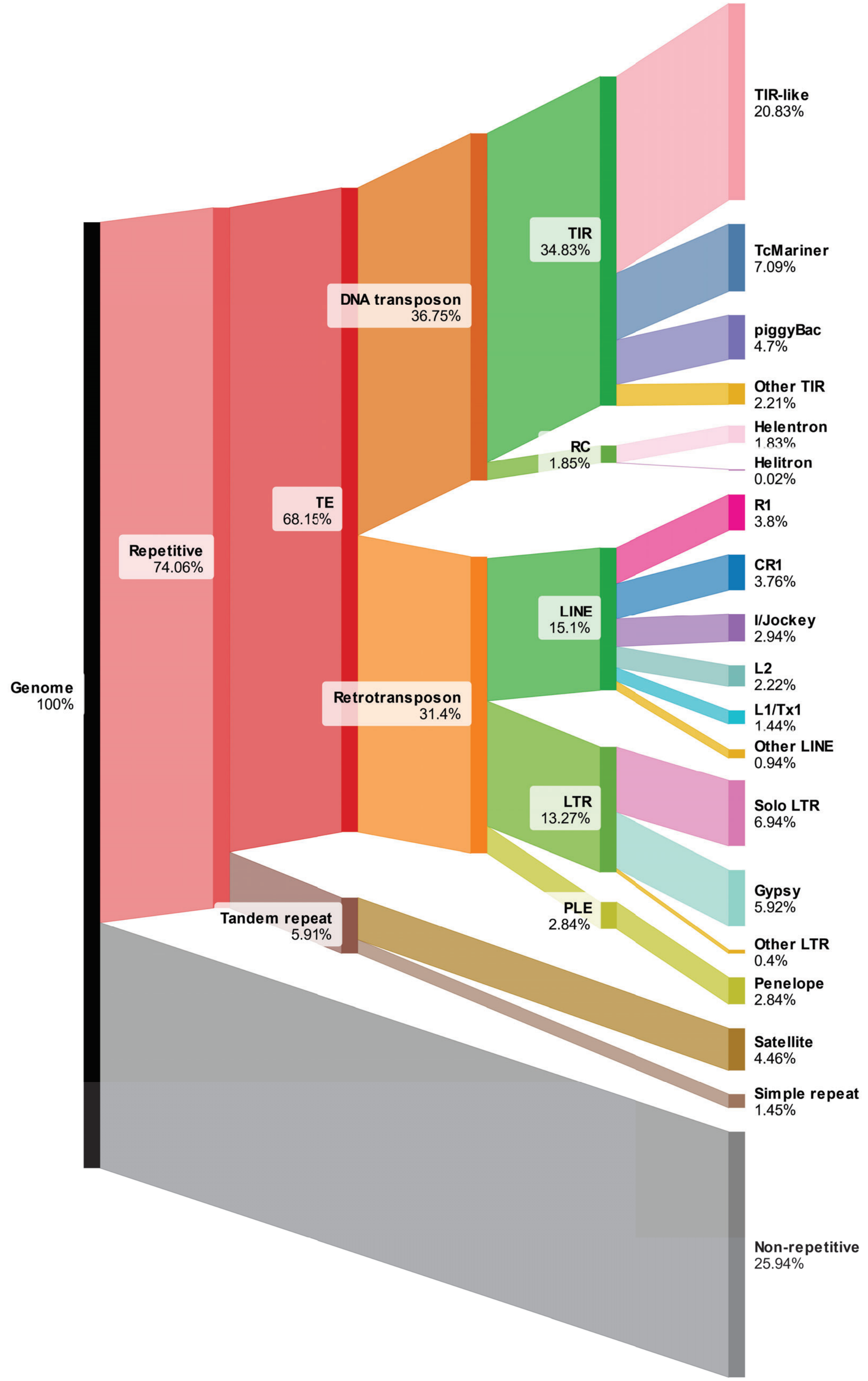
Genome
100%
Repetitive
74.06%
TE
68.15%
Tandem repeat
5.91%
DNA transposon
36.75%
Retrotransposon
31.4%
TIR
34.83%
RC
1.85%
LINE
15.1%
LTR
13.27%
PLE
2.84%
TIR-like
20.83%
TcMariner
7.09%
piggyBac
4.7%
Other TIR
2.21%
Helentron
1.83%
Helitron
0.02%
R1
3.8%
CR1
3.76%
I/Jockey
2.94%
L2
2.22%
L1/Tx1
1.44%
Other LINE
0.94%
Solo LTR
6.94%
Gypsy
5.92%
Other LTR
0.4%
Penelope
2.84%
Satellite
4.46%
Simple repeat
1.45%
Non-repetitive
25.94%

**Figure S6. Genome proportions of repetitive and non-repetitive DNA in (A) *Ixodes ricinus* Murphy and (B) *I. scapularis*.** The majority of the tick genomes are repetitive DNA, dominated by transposable elements (TEs): Terminal Inverted Repeat (TIR) transposons; Long INterspersed Element (LINE) retrotransposons; Long Terminal Repeat (LTR) retrotransposons; Penelope-like (PLE) retrotransposons; and rolling-circle (RC) transposons.

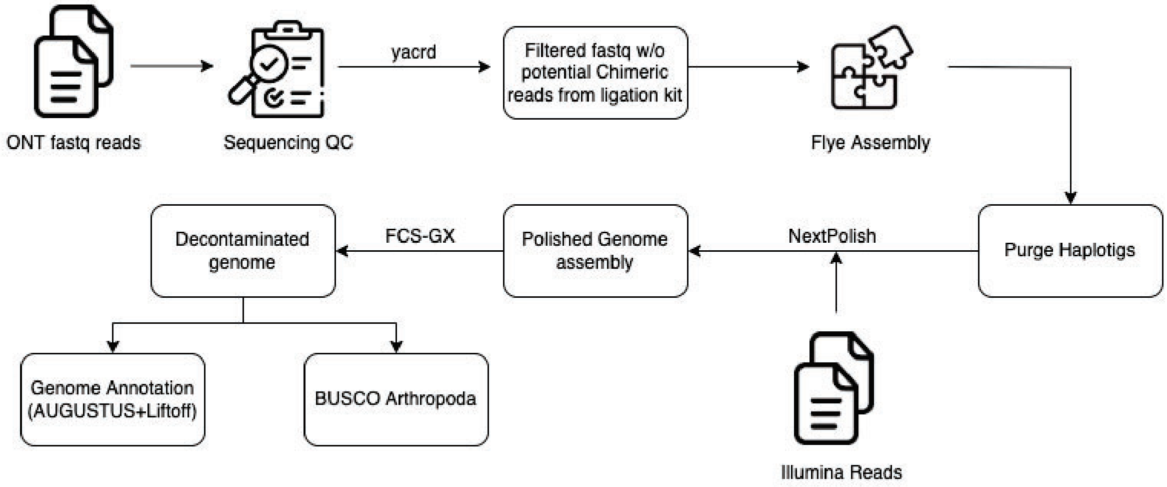


**Figure S7. Genome assembly and analysis pipeline for *Ixodes ricinus*.**

**Table S1. Genome proportions of repetitive and non-repetitive DNA in *Ixodes ricinus* Maya, *I. ricinus* Murphy and *I. scapularis*.** Summary of the *Ixodes* repeat library. Genome proportion percentages were calculated using the total length (bp) reported in Table 1.

**Table S2. Genome assembly of putative scaffolds.** (A) *Ixodes ricinus* Maya and (B) *I. ricinus* Murphy contigs scaffolded to the 14 longest chromosome-scale scaffolds of the current reference genome assembly for *I. scapularis* (NCBI Assembly GCA_016920785.2).

**Supplemental Information Item 1. Randomly selected orthologous intron from *I. ricinus* and *I. scapularis*.** These DNA sequences were used to analyze the accumulation patterns of transposable elements in *Ixodes*.
